# Dietary Zn governs protein: carbohydrate regulation of fecundity and lifespan in *Drosophila melanogaster*

**DOI:** 10.1101/2025.06.27.662062

**Authors:** Sweta Sarmah, Richard Burke, Christen K Mirth, Matthew DW Piper

## Abstract

Recent work has shown that dietary zinc (Zn) restriction has a strong effect to limit egg production in female *Drosophila*, while also producing a variable, but beneficial, effect on lifespan. This combination of phenotypes is interesting because it is consistent with the disposable soma theory of ageing, and phenocopies the well-studied effects of reducing the dietary protein-to-carbohydrate (P:C) ratio. Thus, Zn restriction may be a key precision nutrition intervention to extend lifespan. To investigate this, we decided to assess the interactive effects of dietary Zn and P:C ratios on reproduction and lifespan. We found that the interventions interact, producing equally strong, but not additive, effects to modify female reproduction and lifespan. Specifically, both Zn restriction and lowering the P:C ratio reduced reproduction and benefited lifespan, but only when the other nutrient was maintained at high levels. When either Zn or P:C was at intermediate or low levels, adjusting the other nutritional component varied reproduction, but did so independently of lifespan, which remained fixed at a high level. Disrupting the Zn transporter, *white*, modified the magnitude of some effects, but not their direction. Thus, the micronutrient Zn modifies life-history outcomes to the same extent as altering the macronutrient (P:C) ratio, indicating its nutritional availability is likely to be a key determinant of fitness for *Drosophila*.

## Introduction

Dietary restriction (DR), a well-established intervention that involves reducing calorie intake by 20-50% without causing malnutrition, has been shown to extend health and/or lifespan in a wide range of organisms, from yeast to primates (Holliday 1989, Masoro 2005, Greer 2009, Piper 2011, Fontana 2015). In most cases, DR promotes lifespan extension at the same time as reducing reproduction, which has led to the hypothesis that lifespan is extended by redirecting resources away from reproduction towards self-preserving somatic maintenance systems (Williams 1966, Holliday 1989, Kirkwood 2000). Conversely, when resources are plentiful, it is thought that organisms prioritize their resources for reproduction over somatic maintenance, thus leading to compromised lifespan.

While the traditional view of DR maintained that restricting dietary calories is critical for lifespan extension, recent work has highlighted the importance of specific nutrients, particularly the balance of the macronutrients protein and carbohydrate, in determining lifespan and reproduction (Simpson 2012). In particular, reducing the dietary protein-to-carbohydrate (P:C) ratio can reproduce the inverse relationship between lifespan and reproduction that is observed during DR (Lee 2008, Maklakov 2008, Piper 2008, Skorupa 2008, Fanson 2012, Solon-Biet 2014). Even more recently, alterations in the levels of dietary micronutrients have been shown to play a central role in modifying, and even mediating, the relationship between these life history traits (Zanco 2021, Sarmah 2025)(Kosakamoto 2024). These findings indicate that dietary micronutrient levels may be key determinants of organismal fitness and highlight the need for a more comprehensive understanding of the way they interact with the macronutrients.

In recent work, we and others found that severe dietary Zn restriction reduced reproduction and, in some conditions, also increased lifespan – an effect that is similar to what is seen when dietary P:C is lowered (Simpson 2017, Kosakamoto 2024, Sarmah 2025). These data indicate a positive effect of Zn restriction on lifespan, but in a manner that depends on some unknown aspects of the nutritional context in which it is applied. In their study, Kosakamoto et al. (2024) also showed that severe Zn restriction increases protein appetite in female flies (Kosakamoto 2024). Thus, Zn and protein appear to interact closely to influence organismal behaviours and resource re-allocation priorities.

To investigate this further and understand the priority that *Drosophila* females assign to each nutrient in generating lifespan benefits following diet change, we co-varied the levels of protein and Zn using a synthetic diet. Our data provide new insights into the way nutritional interactions work to modify lifespan, and thus establish a more precise set of nutritional conditions for future studies to characterize the molecular effects of nutrient-sensing pathways and metabolic networks in mediating dietary trade-offs between reproduction and survival.

## Results

### Dietary Zn restriction interacts with protein : carbohydrate changes to determine lifespan and reproduction

To explore the interactive effects of P:C and Zn ratios on lifespan and reproduction, we utilized a completely defined holidic medium, which allowed us to manipulate each nutrient independently (Piper 2014, Sarmah 2024). We tested the effects on lifespan and reproduction of varying Zn levels from that found in our complete diet (100%), to a diet with a limiting amount of Zn (10%), to a diet with no added Zn, which represents extreme restriction (0%; (Sarmah 2025)). We did so for three levels of P:C - low, intermediate and high. To achieve altered P:C ratios, the protein levels were set at 37.5 mM, 75 mM, and 150 mM nitrogen equivalents, with a constant carbohydrate (sucrose) level of 50 mM across all conditions (Figure 1A). These levels were chosen based on previous findings (Wu 2020) that this P:C series associated positively with reproduction, and negatively with lifespan for our outbred *Drosophila* strain Dahomey.

**Figure 1.**
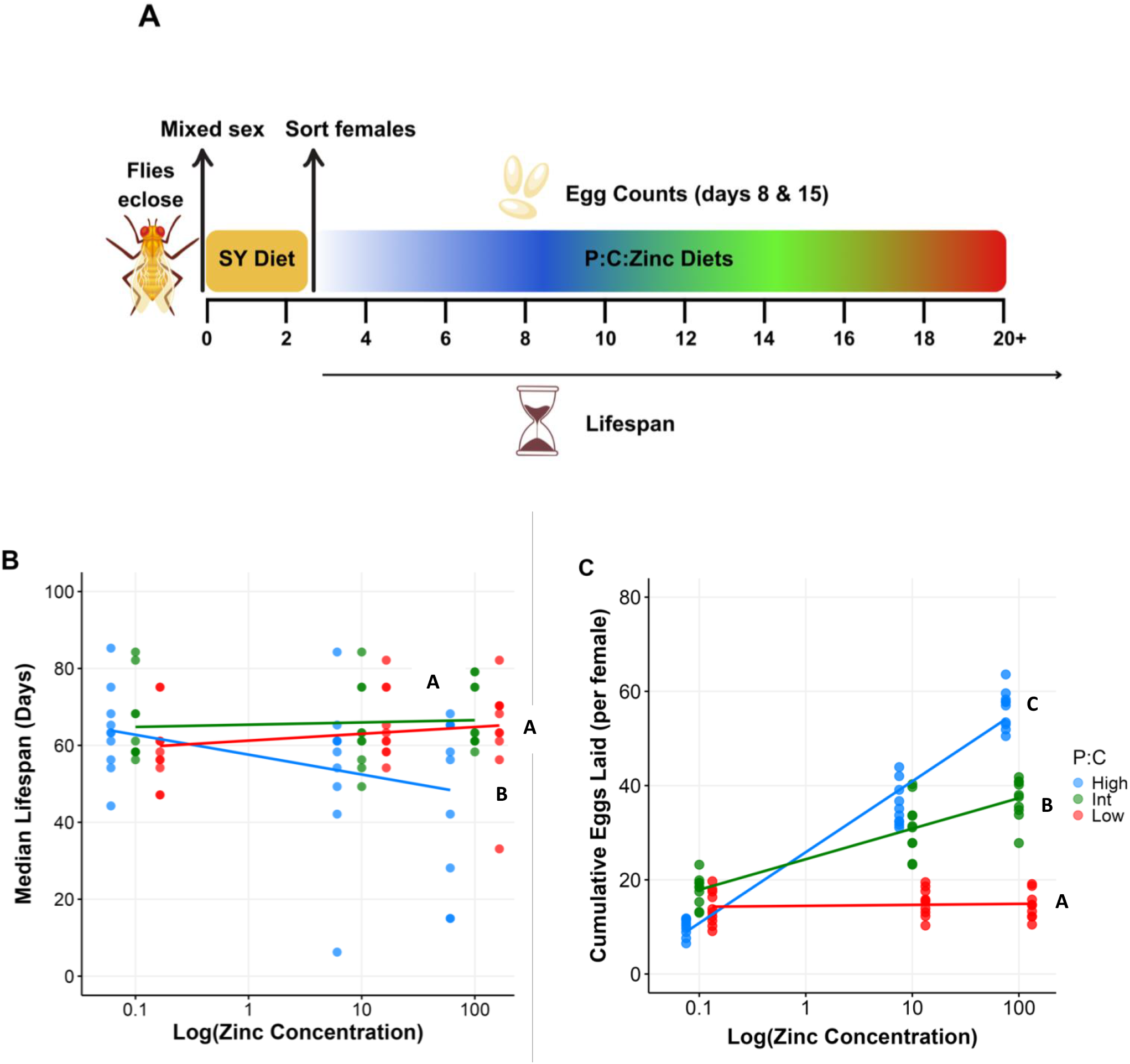
Lifespan and fecundity change in response to Zn restriction in a manner that is modified by dietary P:C concentration. A) Flies were maintained on a sugar-yeast (SY) diet until 2 days of adulthood. On day 3, mated females were isolated and transferred to one of nine synthetic diets containing one of three P:C ratios (low, intermediate, high) at each of three levels of Zn (0%, 10%, 100%). Lifespan was monitored throughout adulthood, and egg production was quantified on days 8 and 15. N = 100 flies per diet (10 flies/vial, 10 vials). B) The response of median lifespan (days) to Zn concentrations (log scale) for each P:C ratio. Zn dilution did not modify lifespan responses to low or medium P:C diets. On a high P:C diet, Zn dilution increases lifespan from a relatively low level. C) Cumulative egg production response to Zn concentrations (log scale) for each P:C ratio. Low P:C diets restricted egg production to a constant level irrespective of Zn concentration. Increasing Zn in the intermediate P:C diet caused fecundity to increase and this effect was further elevated at high P:C. Regression lines with statistically distinct slopes are indicated by different letters (Tables S2 & S4).

For lifespan, we found a significant interaction between the P:C ratio and dietary Zn levels (Figure 1B-C, Figure S1, Table S1). As we previously reported, Zn restriction did not alter lifespan at our intermediate P:C level (Sarmah 2025) (Figure 1B, Figure S1 C, Table S2), and this was no different on the low P:C diets (Figure 1B, Figure S1A, Table S2). By contrast, on a high P:C diet, Zn restriction caused lifespan to increase from a relatively short duration, up to the maximum level observed across all diets tested (Figure 1B, Figure S1 E, Table S2). Together, these data show that Zn restriction, or lowering P:C, can appear to benefit lifespan, but only when diluted in a nutritional background where the other nutrient is at a high level. They also show that high P:C and high Zn levels are not individually detrimental to lifespan, but they are when combined.

Fecundity was also significantly modified by an interaction between P:C ratio and Zn levels (Figure 1C, Table S3). At low P:C, there was no change in egg production with varying Zn levels. However, the slope of the response increased significantly from low to intermediate P:C (0.12 to 4.85) and further again from intermediate to high P:C (4.85 to 11.55) (Figure 1C, Table S4). Thus, egg production responded to both nutrients, exhibiting the appearance of nutritional co-limitation for the Zn and P:C concentrations that we used (Table S3).

In our previous work, we demonstrated a significant impact of the *white* gene, a transporter, on both reproduction and lifespan for flies maintained on different levels of dietary Zn. To assess if *white* mutation also changes the way flies respond to dietary Zn over different P:C levels, we subjected *white* mutant Dahomey to the same set of diets as above. There was no significant genotype effect on the way the diets modified lifespan (Figure S1 B,D,F, Figure S2A, Table S2) but genotype did modify diet effects on fecundity: *white* mutant Dahomey exhibited a positive response to increasing dietary Zn on all three P:C diets, and a blunted increase as P:C levels rose (Figure S2 B, Table S3 – S4). Thus, both genotypes exhibited the same phenotypic trends across all diets, but mutation in the *white* gene significantly altered the strength of the flies’ egg production responses to dietary Zn and P:C levels.

### Zn and P:C both make key contributions to determining a lifespan / reproduction trade-off

Since neither high Zn, nor high P:C alone are detrimental for lifespan, we looked for evidence that their combined negative effects on lifespan could be explained indirectly, via effects on the flies’ fecundity, which would indicate a lifespan / reproduction trade-off. To do this, we investigated the relationship between egg laying and lifespan across all experimental replicates (Figure 2). The slope of lifespan change in response to egg production was no different for flies feeding on variable levels of Zn at low and intermediate P:C ratios, with no evidence of the negative slope expected for a lifespan / reproduction trade-off (Figure 2A-B, Table S6). By contrast, on the high P:C diets only, lifespan was reduced as egg production increased (Figure 2C, Table 6), which is indicative of a lifespan / reproduction trade-off driven by Zn levels. In further experiments, we found that this pattern of change was the same in flies carrying the *white* mutation (Figure S3), but that the slope changes were more exaggerated (Table S6).

**Figure 2.**
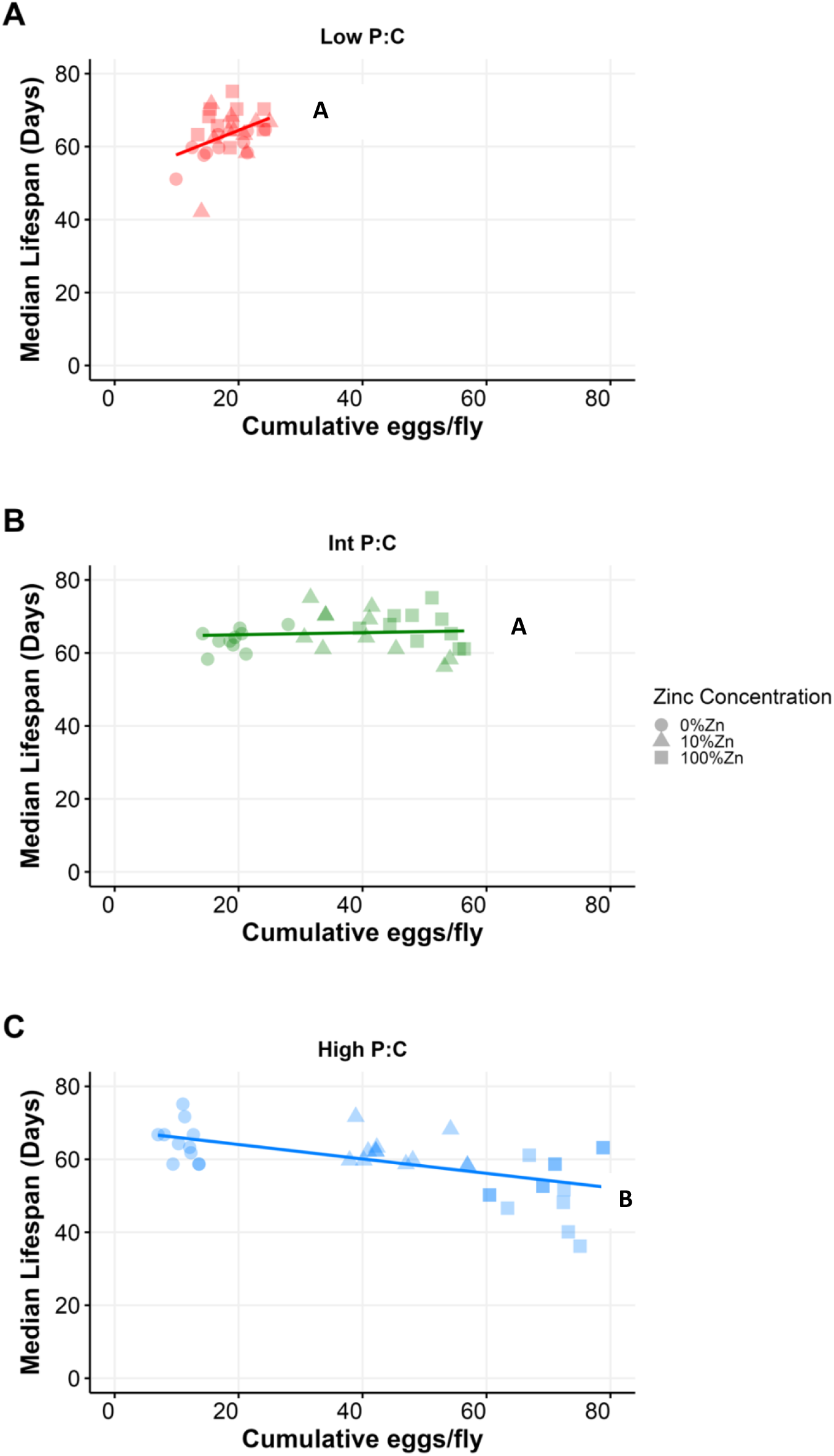
The relationship between cumulative eggs laid per female and median lifespan (days) varies across different P:C ratios as Zn concentration changes. A) The relationship between egg laying and lifespan across Zn concentrations was indistinguishable for flies on low P:C, and B) intermediate P:C diets. C) However, there was a significantly different, negative, relationship between egg laying and lifespan for flies on the high P:C diets, indicating a trade-off. Regression lines with statistically distinct slopes are indicated by different letters (Table S6).

## Discussion

The Disposable Soma theory proposes a mechanism for lifespan extension by DR, in which resource prioritisation is switched between reproduction and somatic maintenance as a response to nutrients being abundant or scarce (Holliday 1989, Kirkwood 1991, Kirkwood 2000, Flatt 2018). The culmination of several years of recent work in *Drosophila* to alter reproduction and lifespan through changes in nutrient balance, rather than food restriction, have produced evidence to both support and refute the mechanism proposed by the disposable soma theory (Grandison 2009, Lee 2014, Piper 2017, Juricic 2020, Fulton 2024, Kosakamoto 2024). For example, when restricting dietary protein during sterol limitation, lifespan is extended and reproduction declines (Zanco 2021), perhaps due to reallocation of sterols between the traits (Zanco 2021). By contrast, supplying flies with small amounts of high-quality protein in a high sterol diet maximises both reproduction and lifespan indicating no trade-off (Zanco 2021). We have shown that Zn restriction can give the appearance of extending lifespan via a reduction in fecundity, but only when the diet contains high P:C levels. Under other conditions in which either Zn or the P:C ratio was maintained at a lower level, fecundity could be varied by changing the level of the other nutrient, but lifespan remained fixed long and unchanged. Together, these data demonstrate that lifespan “extension” in response to Zn restriction or lowered P:C ratios, and the appearance of its trade-off with fecundity, depend on the nutritional context in which they are applied. Further work is required to determine if altering Zn levels in a high P:C diet works through a mechanistic trade-off, as proposed by the disposable soma theory or if high Zn and high P:C are beneficial for egg production but simultaneously toxic for lifespan.

On a technical note, small differences in diet quality can readily explain apparent discrepancies in the relationship between reproduction and lifespan across different labs that use different amounts and proportions of various naturally occurring ingredients when making their fly food (Bass 2007, Piper 2007, Ziehm 2013, Lesperance 2020). Thus, the mechanistic nature of the DR response depends very much on the nutritional context used in each experiment and the degree to which each dietary component is limiting, or close to limiting, in the control diet. It will be interesting in future experiments to combine each of the key limitations we have discussed above, and characterise their molecular interactions in an attempt to gain insights into the mechanisms by which food modifies lifespan.

In mammals, Zn plays a critical role in insulin secretion by stabilizing insulin hexamers in pancreatic β-cell secretory granules, and Zn restriction has been shown to impair insulin production and secretion (Dodson 1998, Chimienti 2006, Wijesekara 2009). Zn also modulates insulin signaling in peripheral tissues to enhance glucose uptake (Norouzi 2018). In *Drosophila*, recent work by Kosakamoto et al. found that dietary Zn restriction extended lifespan, possibly via a mechanism in which Zn-gated chloride channel action is reduced, thus limiting insulin-like peptide (ILP) secretion – an intervention that can extend organismal lifespan by itself (Kosakamoto 2024) (Broughton 2005).Similarly, in *C. elegans*, Zn has been shown to modulate DAF-16/FOXO signalling, a key downstream effector of IIS that regulates stress resistance and lifespan (Kumar 2016). Together, these findings point to the possibility that Zn restriction extends lifespan by reducing IIS activity. It would be interesting to measure the levels of insulin like peptides and IIS target gene expression in flies reared on high P:C diets with varying Zn availability.

In prior work, it has been established that the *white* gene, which encodes a Zn transporter, plays a role in dietary Zn uptake and distribution (Yin 2017, Tejeda-Guzmán 2018), and we found that this gene is important to limit egg production during dietary Zn limitation, thus assuring greater egg viability (Sarmah 2025). Here, we find further evidence for this effect, as *white* mutant flies tended to lay more eggs than red-eyed (*white*+) flies during severe Zn restriction as the P:C ratio increases. We predict that these extra eggs contain a significant proportion that are not viable. In future work, it will be interesting to determine the tissue specific role of *white* to limit egg production in response to low Zn levels.

In natural environments, Zn is often limiting, with its availability in fruit heavily dependent on soil content, a factor that varies geographically and seasonally (Alloway 2009). Yeast, a key component of *Drosophila* diets, dynamically regulates Zn homeostasis through specialized transporters and Zn-sensing transcription factors (Wilson 2016). Together, these factors could alter Zn bioavailability to flies feeding on fermenting fruit. Given that Zn is essential for reproductive success (Hu 2020, Sarmah 2025), organisms may utilise Zn levels as an indicator of environmental quality to determine their reproductive strategy. In fact, we found here that Zn has a role that is equally as important as the macronutrients since Zn restriction was just as effective as protein restriction to limit female egg production. In *C. elegans*, Zn-limited conditions have been shown to alter oogenesis and decrease fertility in hermaphrodites (Hester 2017), while in humans, Zn deficiency can occur at high levels in regions relying heavily on cereal-based diets with low Zn content caused by poor soil quality (Wessells 2012)(Cakmak 2008). The effects of Zn deficiency include delayed sexual maturation, miscarriage, infertility, and pregnancy complications (Goldenberg 1995, Shah 2001, Prasad 2013, Nossier 2015). Thus, the effects of Zn restriction on reproduction are widespread, pointing to the general importance of this dietary micronutrient in regulating fitness outcomes.

## Methods

### Fly husbandry

All experiments used outbred “wild-type” *Drosophila* of the red-Dahomey (*rDah*) strain. Stocks were maintained in large population cages at 25°C under a 12-hour light/dark cycle, ensuring continuous overlapping generations. Prior to experimentation, flies were removed from the population cages and reared for two generations at controlled densities, using eggs laid by age-matched mothers to minimize parental effects (Linford 2013). Once the third generation emerged, newly eclosed adults mated for 48 hours before being lightly anesthetized with CO2 and sorted by sex. Throughout, *rDah* stocks were maintained on sugar-yeast (SY) food (Bass 2007).

### Experimental diets (Holidic medium)

To examine how varying the P:C:Zn ratio affects lifespan and fecundity of female fruit flies, we prepared a series of holidic diets following protocols adapted from *Piper et al*. (Piper 2014). First, we generated three levels of total amino acids (37.5 mM, 75 mM, and 150 mM), converted to protein equivalents using the nitrogen-based calculation (assuming that nitrogen constitutes 16% of the protein; (Sosulski 1990)). The amino acid composition was matched to the adult *Drosophila* exome (FLYAA) ratio as described by *Piper et al*. (Piper 2017). All diets contained a constant carbohydrate source (sucrose) at 50 mM. Three Zn levels (100%, 10%, or 0%) were used of the normal level used in standard holidic medium. Each of these dietary combinations was formulated to yield comparable total caloric densities, thereby isolating the effects of the altered protein and Zn levels. As with previous work demonstrating that varying amino acid and micronutrient levels affects key life-history traits (Piper 2017, Wu 2020), these diets allowed us to systematically dissect the contributions of protein, carbohydrate, and Zn to both lifespan and reproductive output in female fruit flies.

### Lifespan assay

For each experimental diet, 10 female flies were placed into each of 10 vials (10 replicates). Every 2-3 days, flies were transferred into new vials containing fresh food at which point deaths and censors were recorded (Piper 2016) and saved using the software Dlife (Linford 2013).

### Fecundity assays

For the fecundity assay, a web camera mounted on a Zeiss dissecting microscope was used to capture digital images of the eggs on the food surface. Egg counts were performed manually from these images. Egg production was measured on days 8 and 15 of the experiment, following 24 hours of dietary exposure. Fecundity was reported as the number of eggs laid per female during each laying period. Each diet treatment consisted of 10 vials, with 10 flies per vial.

### Statistical Analyses

Statistical analyses were performed using R version 4.3.1 (2023-06-16). Linear mixed-effects models (LMMs) and linear models (LMs) were implemented with the lme4 (Bates 2015) and lmerTest packages. Significance of fixed effects was evaluated using Type III Wald chi-square tests from the car package (Fox 2019). Post hoc comparisons were conducted with emmeans (Lenth 2021) and adjusted using Tukey’s method. Plots were produced using ggplot2 (Wickham 2016).

Median lifespan (AgeD) was analysed using LMMs with Zn concentrations (log transformed), P:C ratio, genotype, and their interactions as fixed effects, and Vial as a random effect. Model simplification followed a stepwise approach: non-significant higher-order interactions (e.g., the three-way log (Zn): Genotype : PC interaction, *p* = 0.348) were removed if likelihood ratio tests (LRTs) and AIC comparisons did not reveal a significant worsening in model fit. Slopes of Zn effects across P:C levels were estimated using emtrends, with pairwise differences assessed via Tukey’s tests (Table S2).

Like median lifespan, egg production was modelled with LMMs, including Treatment (P:C ratio), Genotype, log10(Zinc+1) (to accommodate 0 values), and their interactions as fixed effects, with Vial as a random effect. Slopes for Zn effects were compared within genotypes using Tukey-adjusted pairwise tests.

The association between cumulative egg production and median lifespan was analyzed with LMs stratified by genotype. Models included Treatment (P:C-Zn), egg count, genotype, and their interaction as predictors. Significant interactions (e.g., Treatment : Eggs : Genotype, p < 0.05; Table S5) prompted slope comparisons across P:C levels using emtrends.

## Author Contributions

Sweta Sarmah: conceptualisation, data curation, formal analysis, investigation, methodology, visualisation, writing – original draft. Richard Burke: conceptualisation, formal analysis, funding acquisition, methodology, supervision, and writing – review and editing Christen K. Mirth: conceptualisation, formal analysis, funding acquisition, methodology, supervision and writing – review and editing. Matthew D. W. Piper: conceptualisation, funding acquisition, methodology, resources, supervision, and writing – review and editing.

## Data Availability Statement

The data that support of the findings of this study are available in Figshare (DOI: 10.26180/29424596)

