## Supplemental Table for "Dietary Zn governs protein: carbohydrate regulation of fecundity and lifespan in *Drosophila melanogaster*"

Table S1.
Analysis of Deviance (Type III Wald Chi-square Tests) of the generalized linear mixed model to best represent the relationship between dietary P:C levels, Zn concentration, and genotypes (red-eyed and white eyed Dahomey) and their interactions on median lifespan. Zn concentrations were natural log-transformed (ln).

| | Chisq | Df | *P value* |
|---|---|---|---|
| ln(Zn) | 84.83 | 1 | <0.001*** |
| Genotype | 0.74 | 1 | 0.3889 |
| P:C | 9.39 | 2 | <0.01** |
| ln(Zn): Genotype | 1.70 | 1 | 0.1925 |
| ln(Zn) : P:C | 116.85 | 2 | <0.001*** |

Table S2.
Differences in the way median lifespan responded to dietary Zn level across different P:C ratios were assessed using Tukey's multiple comparisons tests (emtrends). Treatment slopes that share a common letter (uppercase for red-eyed Dahomey, lowercase for white eyed Dahomey ) are statistically indistinguishable.

Genotype= *rDah*

| | Treatment. trend | Df | group |
|---|---|---|---|
| Low P:C | 0.881 | 82 | A |
| Intermediate P:C | -0.062 | 98 | A |
| High P:C | -2.333 | 71 | B |

Genotype= *wDah*

| | Treatment. trend | Df | group |
|---|---|---|---|
| Low P:C | 0.965 | 90 | a |
| Intermediate P:C | -0.313 | 85 | a |
| High P:C | -3.254 | 80 | b |

Table S3.
Analysis of Deviance (Type III Wald Chi-square Tests) of the generalized linear mixed model to represent relationship between genotypes (*rDah* & *wDah*), P:C treatment and dietary Zn levels on fecundity. Zn concentrations were log10-transformed after adding 1 to accommodate zero values.

| | Chisq | Df | *P value* |
|---|---|---|---|
| P:C Treatment | 4.62 | 2 | 0.099 |

| | | | |
|---|---|---|---|
| Genotype | 4.98 | 1 | <0.05* |
| Log10(Zn+1) | 122.88 | 1 | <0.001*** |
| P:C Treatment: Genotype | 2.59 | 2 | 0.274 |
| P:C Treatment: log10(Zn+1) | 60.77 | 2 | <0.001*** |
| Genotype: log10(Zn+1) | 0.78 | 1 | 0.376 |
| P:C Treatment: Genotype: log10(Zn+1) | 8.06 | 2 | <0.05* |

Table S4.
Differences in fecundity across different P:C ratios and Zn concentrations for genotypes (*rDah* & *wDah*), measured using Tukey's multiple comparisons tests (emtrends).

Genotype= *rDah*

| Treatment | Log10(Zn+1). trend | Df | group | *P value* |
|---|---|---|---|---|
| Low P:C | 0.12 | 84 | A | 0.908 |
| Intermediate P:C | 4.85 | 84 | B | <0.001*** |
| High P:C | 11.55 | 84 | C | <0.001*** |

Genotype= *wDah*

| Treatment | Log10(Zn+1). trend | Df | group | *P value* |
|---|---|---|---|---|
| Low P:C | 2.78 | 168 | a | <0.05* |
| Intermediate P:C | 9.33 | 168 | b | <0.001*** |
| High P:C | 10.24 | 168 | b | <0.001*** |

Table S5.
Anova (type III) of the linear model to assess the effects of dietary treatments (Zn : P : C) and genotype (red-eyed and white eyed Dahomey) on the relationship between egg production and median lifespan.

| | Sum Sq | Df | *P value* |
|---|---|---|---|
| Treatment | 311 | 2 | <0.01** |
| Eggs | 880 | 1 | <0.001*** |
| Genotype | 0 | 1 | 0.942 |
| Treatment: Eggs | 521 | 2 | <0.01** |
| Treatment: Genotype | 75 | 2 | 0.309 |
| Eggs: Genotype | 441 | 1 | <0.001*** |
| Treatment: Eggs: Genotype | 237 | 2 | <0.05* |

Table S6.
Variations in the relationship between egg production and lifespan across the three P:C ratios using data from the three Zn levels combined, stratified by genotype (red-eyed and white-eyed Dahomey). Differences in slope were measured using Tukey's multiple comparisons tests (emtrends).

Genotype= *rDah*

| Treatment | eggs. trend | Df | group |
|---|---|---|---|
| Low P:C | 0.669 | 92 | A |
| Intermediate P:C | 0.029 | 92 | A |
| High P:C | -0.197 | 92 | B |

Genotype= *wDah*

| Treatment | eggs. trend | Df | group |
|---|---|---|---|
| Low P:C | 0.672 | 92 | a |
| Intermediate P:C | 0.089 | 92 | b |
| High P:C | -0.421 | 92 | c |
