## Supplemental Figures for "Dietary Zn governs protein: carbohydrate regulation of fecundity and lifespan in *Drosophila melanogaster*"

**Figure S1**

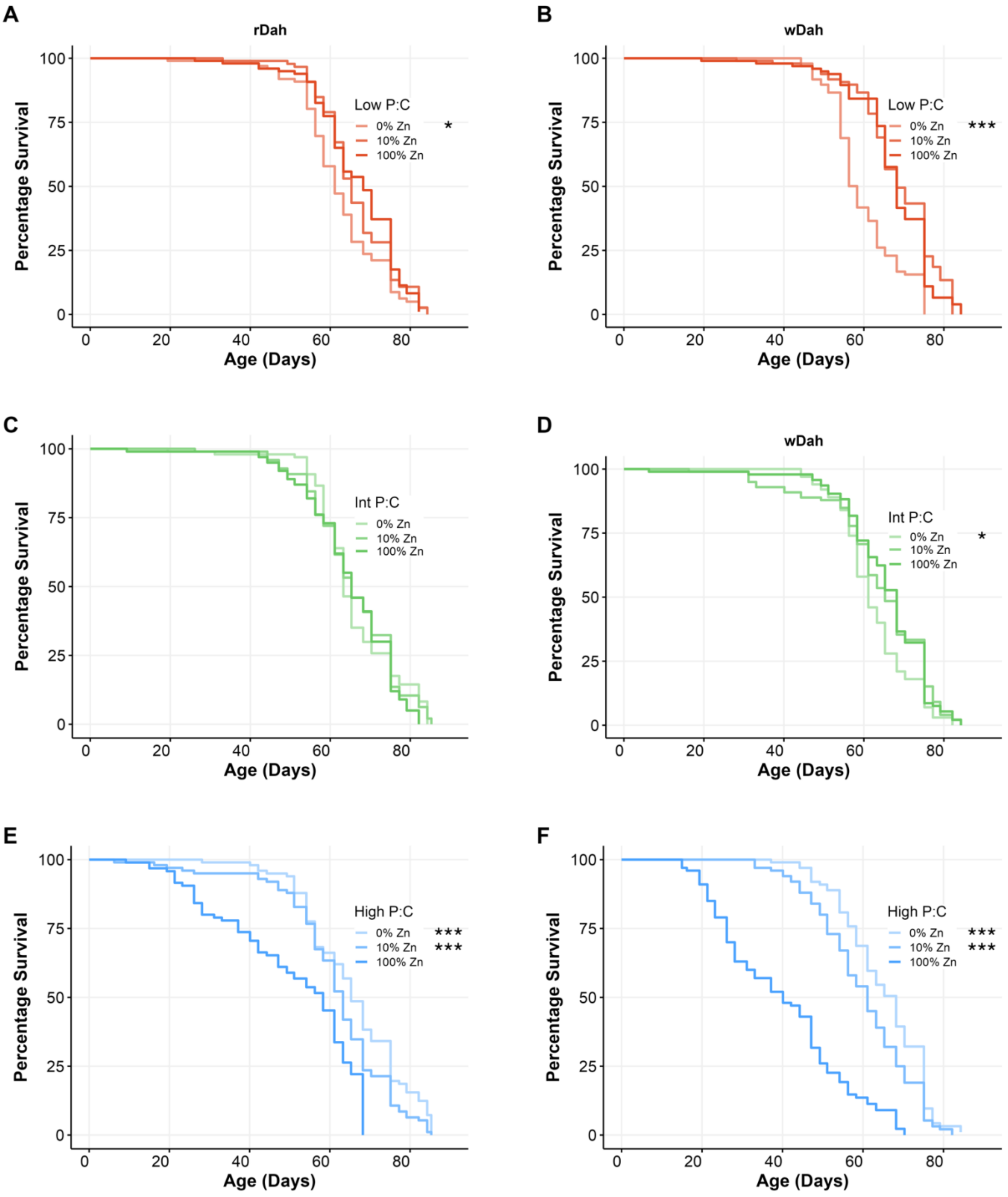
A
rDah
Percentage Survival
100
75
50
25
0
Low P:C
0% Zn
10% Zn
100% Zn
*
Age (Days)
0
20
40
60
80
B
wDah
Low P:C
0% Zn
10% Zn
100% Zn
***
C
Int P:C
0% Zn
10% Zn
100% Zn
D
wDah
Int P:C
0% Zn
10% Zn
100% Zn
*
E
High P:C
0% Zn
10% Zn
100% Zn
***
***
F
High P:C
0% Zn
10% Zn
100% Zn
***
***

***Figure S1. Survival curves of adult rDah and wDah female flies maintained on different P:C and Zn concentrations (raw data for that shown in Figure 1)***

*(A-B) Low P:C diet: compared with 100% and 10% Zn, which were indistinguishable, rDah showed a small but significant decrease in lifespan at 0% Zn ( $p=0.015$), which was exaggerated for wDah ( $p<0.001$).(C-D) Intermediate P:C diet: No significant effects on lifespan were observed in rDah ($p>0.05$), but wDah showed a small but significant decrease in lifespan at 0% Zn ( $p=0.011$). (E-F) High P:C diet: rDah showed a significant increase in lifespan with Zn restriction to 10% or 0% ($p<0.001$), while wDah displayed a significant increase in lifespan from 100% to 10% Zn ( $p<0.001$) and again from 10% to 0% Zn ($p<0.001$).*

**Figure S2**

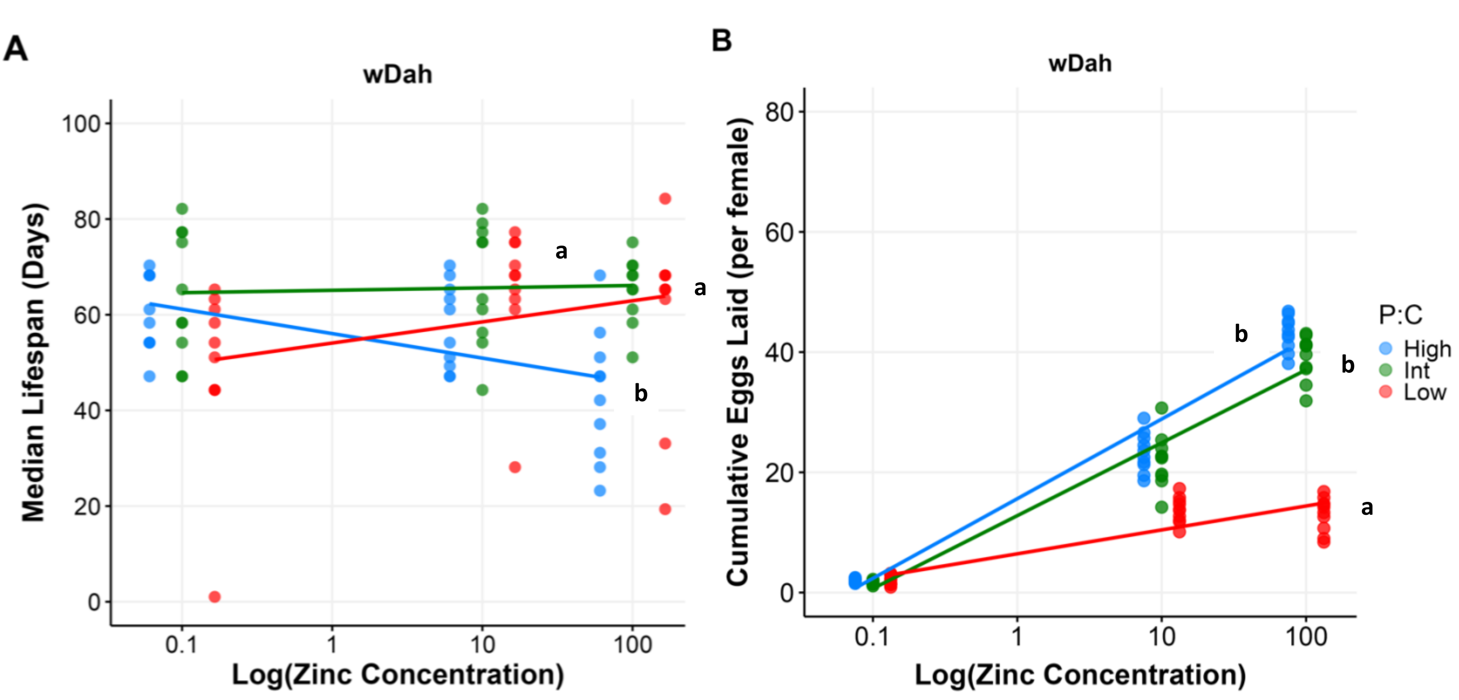


***Figure S2. Lifespan and fecundity responses to Zn restriction in the wDah genotype, modified by dietary P:C concentration***

A) *Flies were maintained on a sugar-yeast (SY) diet until 2 days of adulthood. On day 3, mated females were transferred to one of nine synthetic diets containing one of three P:C ratios at each of three levels of Zn. Lifespan was monitored throughout adulthood, and egg production was quantified on days 8 and 15. N = 100 flies per diet (10 flies/vial, 10 vials).*

B) *The response of median lifespan (days) to Zn concentrations (log scale) for each P:C ratio. At low P:C, increasing Zn significantly extended lifespan, at intermediate P:C, Zn had no effect, while at high P:C, higher Zn shortened lifespan*

C) *Cumulative egg production response to Zn concentrations (log scale) for each P:C ratio. At low P:C, fecundity showed a weak positive association with Zn. At intermediate and high P:C, fecundity increased more strongly with Zn.*

*Regression lines with statistically distinct slopes are indicated by different letters (Tukey's test; Tables S2, S4).*

**Figure S3**

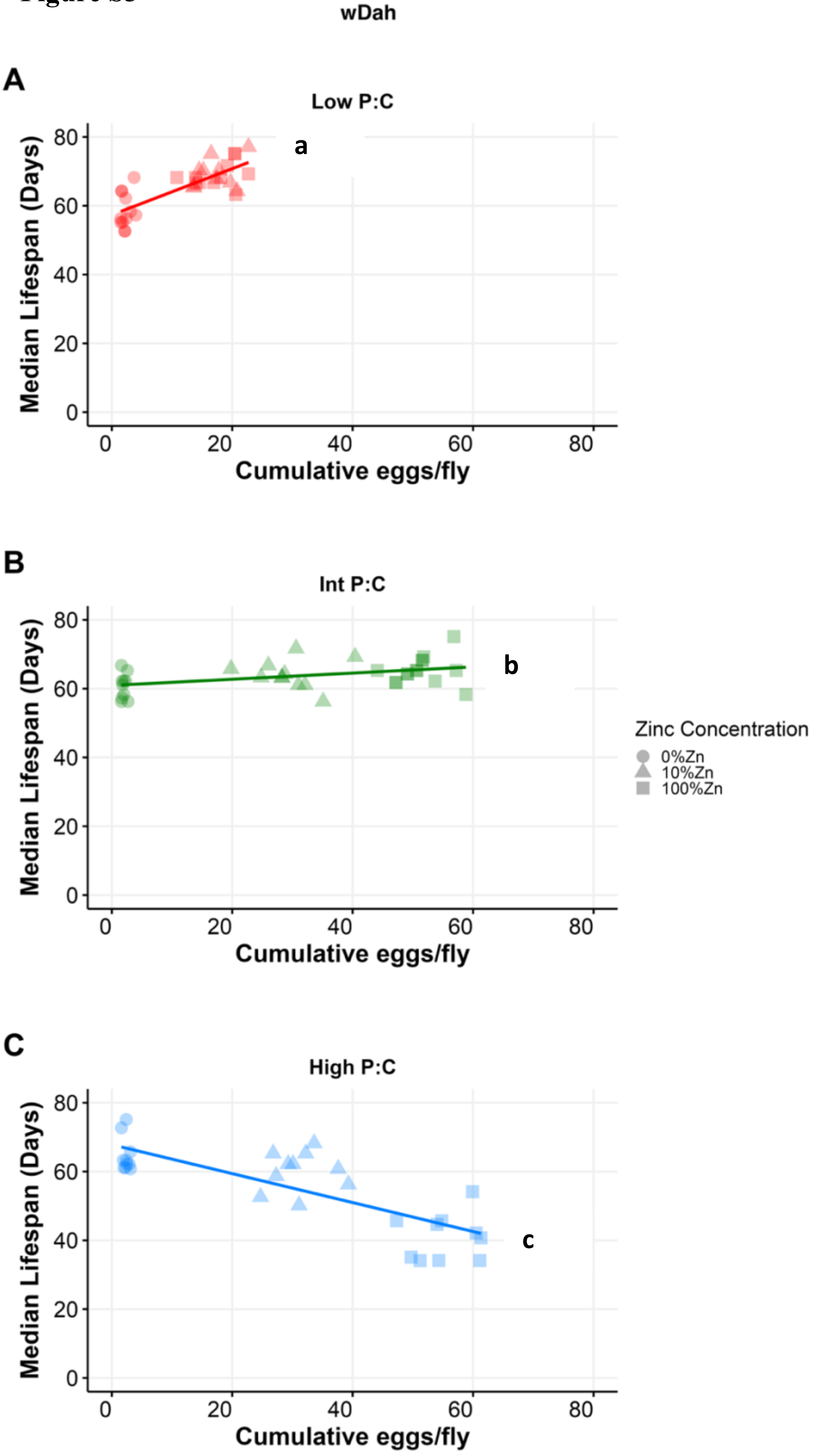

***Figure S3. Relationship between cumulative eggs laid per female and median lifespan (days) in wDah flies across P:C ratios and Zn concentrations***

*A) At low P:C, higher egg production was associated with extended lifespan, indicating synergistic effects between reproduction and survival.*

*B) At intermediate P:C, the relationship shifts to a weak positive association, suggesting that egg production and lifespan are not in direct conflict under these nutritional conditions.*

*C) At high P:C, increased egg production corresponded to sharply reduced lifespan, reflecting a trade-off between reproduction and survival.*

*Regression lines with statistically distinct slopes are indicated by different letters (Table S6).*
